# Biologically realistic mean-field models of conductance-based networks of spiking neurons with adaptation

**DOI:** 10.1101/352393

**Authors:** Matteo di Volo, Alberto Romagnoni, Cristiano Capone, Alain Destexhe

## Abstract

Accurate population models are needed to build very large scale neural models, but their derivation is difficult for realistic networks of neurons, in particular when nonlinear properties are involved such as conductance-based interactions and spike-frequency adaptation. Here, we consider such models based on networks of Adaptive exponential Integrate and fire excitatory and inhibitory neurons. Using a Master Equation formalism, we derive a mean-field model of such networks and compare it to the full network dynamics. The mean-field model is capable to correctly predict the average spontaneous activity levels in asynchronous irregular regimes similar to in vivo activity. It also captures the transient temporal response of the network to complex external inputs. Finally, the mean-field model is also able to quantitatively describe regimes where high and low activity states alternate (UP-DOWN state dynamics), leading to slow oscillations. We conclude that such mean-field models are “biologically realistic” in the sense that they can capture both spontaneous and evoked activity, and they naturally appear as candidates to build very large scale models involving multiple brain areas.

## 1 Introduction

Large-scale models of the brain can be built at cellular resolution (Markram et al., 2015), but this approach requires huge computational resources. Another approach is to build models where the smallest unit is not a neuron, but is a population of neurons, which corresponds to the resolution in imaging studies. Several examples of such a mesoscopic approach have been proposed (reviewed in (Sanz Leon et al., 2013; Deco et al., 2015; Breakspear, 2017; Bassett et al., 2018)). However, such models use representations of neural populations which are mostly phenomenological and often use linear models, and are thus non-realistic because they miss essential non-linear effects, such as conductance-based interactions, or adaptation dynamics.

In the present paper, we would like to propose a first step towards a “biologically realistic” mesoscopic model of neural populations by explicitly including non-linear effects. We use a mean-field approach based on a Master Equation formalism describing the dynamics of spiking neurons (El Boustani and Destexhe, 2009), which we modify so that it can account for both conductance-based interactions and spike-frequency adaptation, yielding a population model which we compare to the cellular-level model.

To be biologically realistic, we focus on several essential features. First, cerebral cortex has a high level of spontaneous activity in the adult mammalian brain. The dynamical regimes observed experimentally in cerebral cortex range from asynchronous states, typically in wakefulness, to regimes displaying slow oscillations consisting of alternating high and low activity states (UP and DOWN states), typically in slow-wave sleep (Dehghani et al., 2016; Renart et al., 2010; Sanchez-Vives and McCormick, 2000; Jercog et al., 2017; Sanchez-Vives and Mattia, 2014; Capone et al., 2017). These states have a common ground of an irregular spiking activity of single neurons (Steriade et al., 2001), while their interaction is known to be mediated by conductance-based synapses (Destexhe et al., 2003). The role of irregularity in neurons activity has been proposed to be important for neurons responsiveness and learning (Denève and Machens, 2016). Because of this feature, the typical asynchronous state observed during awake animals recording is usually named as asynchronous irregular (AI).

A second essential feature is the presence of conductances, and their associated nonlinearity. Conductances have been observed to play a key role in network responses to external input, as different states of the system can lead to different outputs to the same specific stimuli (Zerlaut and Destexhe, 2017). Accordingly, these features should be taken into account in a realistic model of cortical populations. At the cellular level, several spiking network models were proposed, including conductance-based interactions (for example see (Vogels and Abbott, 2005; Destexhe, 2009)). Such models typically use the classic integrate-and-fire (IF) model, and they can reproduce different dynamical states, such as AI states and UP/DOWN states.

A third important feature is the presence of spike-frequency adaptation. This form of adaptation is present in virtually all excitatory neurons in cerebral cortex, which typically display slowly adapting spike frequency responses, a pattern which was called “regular spiking” (RS), in contrast to inhibitory neurons which often fire at higher frequencies with no adaptation, which was called “fast spiking” (FS) neurons (Connors and Gutnick, 1990). Such patterns can be modeled using Hodgkin-Huxley models (Pospischil et al., 2008), or by IF models augmented with the mechanisms allowing spike-frequency adaptation. One of the simplest of such models are two-dimensional IF models including an adaptation variable (Izhikevich, 2003) or the Adaptive Exponential (AdEx) IF model (Brette and Gerstner, 2005). The AdEx model is able to simulate the main cell types in the thalamo-cortical system, such as RS and FS neurons, as well as various types of bursting neurons such as that found in the thalamus (Destexhe, 2009).

Several derivations of mean-field models of networks of spiking neurons have been proposed, mostly using current-based (CUBA) interactions (for example, see (Renart et al., 2003)). Using the IF model, it is possible to approximate the neuron transfer function (TF) (i.e. the output firing rate of a single neuron in function of its inputs). Phenomenological approaches, based on the assumption of a linear transfer function, have been used (Shriki et al., 2003) in order to reproduce the rate response of the network. More recently, mean-field models were successful to reproduce network dynamics (Augustin et al., 2017; Schwalger et al., 2017; Montbrió et al., 2015), but such models cannot account for the presence of non-linearities such as conductances or spike-frequency adaptation. A formalism based on a Master Equation approach was proposed (El Boustani and Destexhe, 2009), and is general enough to predict both average population rates and their covariances, for both current-based and conductance-based networks. Such a formalism, however, requires that the neuronal TF is known analytically, which is not the case for non-linear neurons.

A significant advance was realized recently by proposing a semi-analytical approach (Zerlaut et al., 2016, 2018) for the calculation of the TF of AdEx neurons, which could be potentially applicable to any neuron model and, more interestingly, to biological neurons. This approach permits to build a mean-field model of AdEx networks (Zerlaut et al., 2018). This model can predict network responses to some extent, but cannot account for the effects due to adaptation, which is manifested in the time course of the network response, or the response to oscillatory inputs which is poorly captured.

As these features are essential to obtain a realistic population model, we propose here a mean-field model that include these effects. We restart from first principles, and include adaptation in the Master Equation approach, leading to a mean-field model of spiking networks including adaptation. We then compare this new formalism to a network model of RS-FS AdEx neurons, and in particular focusing on the network transient response to complex stimuli.

## 2 Model and mean field derivation

We describe in this section the spiking network model and the derivation of the corresponding mean-field equations.

### 2.1 Spiking network models

We consider a population of *N* = 10^4^ neurons connected over a random directed network where the probability of connection between two neurons is *p* = 5%. We consider excitatory and inhibitory neurons, with the 20% inhibitory neurons. The dynamics of each of the two types of neurons is based on the adaptive integrate and fire model described by the following equations

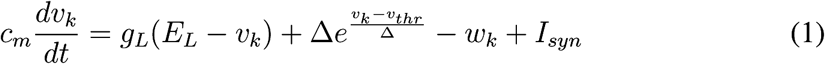

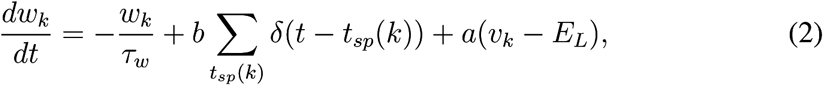

where *c*_*m*_ = 150pF is the membrane capacity, *v*_*k*_ is the voltage of neuron *k* and, whenever *v*_*k*_*> v*_*thr*_ = 50*mV* at time *t*_*sp*_(*k*), *v*_*k*_ is reset to the resting voltage *v*_*rest*_ = 65*mV* and fixed to that value for a refractory time *T*_*refr*_ = 5*ms*. The leak term has a fixed conductance of 10nS and the leakage reversal *E*_*L*_ is varied in our simulations but is typically -65mV. The exponential term has a different strength for RS and FS cells, i.e. Δ = 2mV (Δ = 0.5mV) for excitatory (inhibitory) cells. Inhibitory neurons are modeled according to physiological insights as fast spiking FS neurons with no adaptation while excitatory regular spiking RS neurons have a lower level of excitability due to the presence of adaptation (while *b* varies in our simulations we fix *a* = 4nS and *τ*_*w*_ = 500ms if not stated otherwise). The synaptic current *I*_*syn*_ received by neuron *i* is the result of the spiking activity of all pre-synaptic neurons *j* ∈ pre(*i*) of neuron *i*. This current can be decomposed in the result received from excitatory E and inhibitory I pre-synaptic spikes 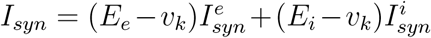, where *E*_*e*_ = 0mV (*E*_*i*_ = 80mV) is the excitatory (inhibitory) reversal potential. Notice that we consider voltage dependent conductances. Finally, we model 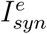 as an decaying exponential function that takes kicks of amount *Q*_*E*_ at each pre-synaptic spike, i.e.:

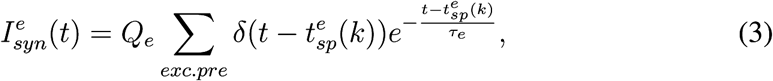

where *τ* _*e*_ = *τ*_*i*_ = 5ms is the decay time scale of excitatory and inhibitory synapses and *Q*_*e*_ = 1nS (*Q*_*i*_ = 5nS) the excitatory (inhibitory) quantal conductance. We will have the same equation with *E* → *I* for inhibitory neurons.

### 2.2 Mean-field equations

For the theoretical analysis of asynchronous dynamics in sparsely connected random networks we use the mean-field approach proposed by El Boustani & Destexhe in (El Boustani and Destexhe, 2009). In this framework, based on a master equation formalism, differential equations for the population average firing rate *v*_*e*_ (*v* _*i*_) of the excitatory (inhibitory) populations in the network are derived, with a time resolution *T*. In fact, the main hypothesis in this formalism, consists in considering the system memoryless after a time *T*, or, in other words, to consider a Markovian dynamics for the network.

In this paper we extend the model described in (El Boustani and Destexhe, 2009) by including the effects of adaptation. The same formalism indeed easily allows this kind of extension, as far as the time scale that drives the dynamics of the adaptation variable *w*(*t*) is slow with respect to the time scale of the mean-field formalism *T*. It is possible to show that when a generic slow variable *W* (with time constant much longer than the mean-field time-scale *T*) is added to the system, one can close the equations for the population activities following (El Boustani and Destexhe, 2009) considering *X* as stationary at every step of the Makrovian process *T* (see Appendix 3.5). When applied to the simpler case discussed in this paper of one excitatory and one inhibitory populations, with population activities *µ* = {*e,i*}, the differential equations read:

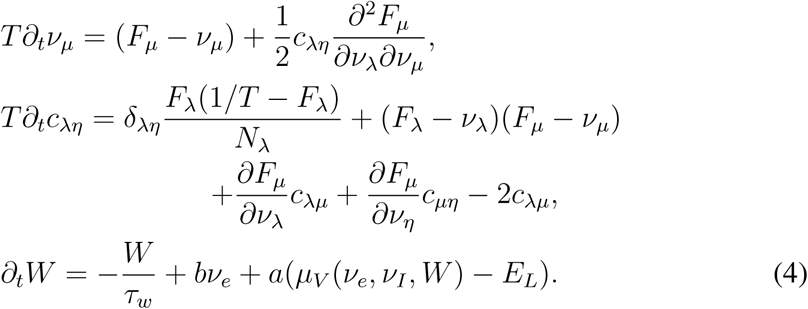

where *F*_*µ*={*e,i*}_ = *F*_*µ*={*e,i*}_(*v*_*e*_, *v*_*i*_,*W*) is the transfer function of a neuron of type *µ*, i.e. its output firing rate when receiving excitatory and inhibitory inputs with rates *v*_*e*_ and *v*_*e*_ and with a level of adaptation *W*. Accordingly here the transfer function is now a function not only of the firing rate *v*_*e*_ and *v*_*i*_, but also of the adaptation *W*. The calculation of the average population voltage *µ*_*V*_ (*v*_*e*_, *v*_*I*_,*W*) is described in the following sections.

### 2.3 Neurons transfer function

We perform a semi-analytical derivation of the transfer function *TF* of RS and FS neurons following (Zerlaut et al., 2016). Here we take explicitly into account the effect of adaptation for the transfer function calculation as the firing rate of a single neuron depends on the input firing rate but also on the adaptation current *w* affecting its voltage dynamics. The method is based on the hypothesis that the output firing rate of a neuron can be written as a function of the statistics of its sub-threshold voltage dynamics, i.e. the average sub-threshold voltage *µ*_*V*_, its standard deviation *μ*_*V*_ and its time correlation decay time *τ*_*V*_.

We report first how to evaluate (*µ*_*V*_, *σ*_*V*_, *τ*_*V*_) as a function of the inputs firing rates (*v*_*E*_,*v*_*I*_) and the adaptation intensity *w*.

### 2.1 From input rates to sub-threshold voltage moments

Following (Kuhn2004), the mean membrane potential is obtained by taking the stationary solution to static conductances given by the mean synaptic bombardment with firing rates (*v*_*E*_, *v*_*I*_). We can calculate the average *µ*_*Ge,Gi*_ and standard deviation _*Ge,Gi*_ of such bombardment for both excitatory and inhibitory process (described by Eq. (3)) in the case spikes follow a Poissonian statistics:

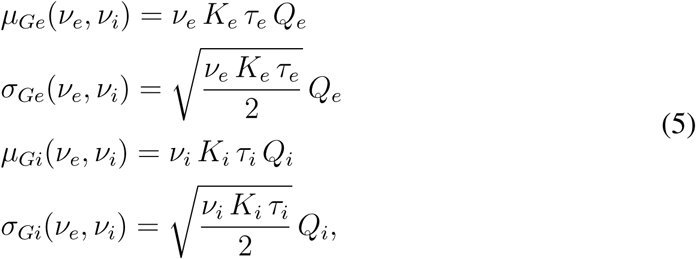

The mean conductances will control the input conductance of the neuron *µ*_*G*_ and therefore its effective membrane time constant *τ*_*m*_:

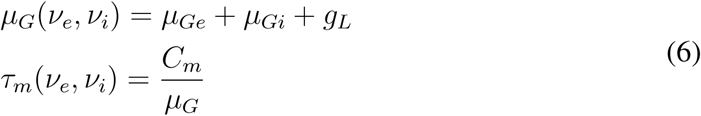

For a specific value *w* of the adaptation current (whose dynamics we remember to be much slower that voltage fluctuations) we obtain the following formula for the average voltage (neglecting the exponential term in Eq.(1)):

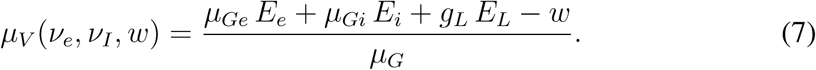

The calculation of σ_*V*_ and of *τ*_*V*_ is identical to (Zerlaut et al., 2018), we just report the final formula:

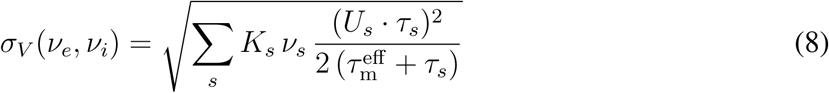

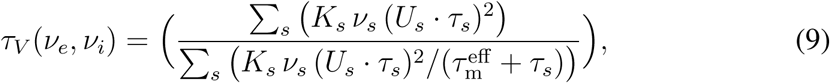

where we defined 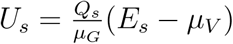.

### 2.2 From sub-threshold voltage moments to the output firing rate

Once calculated (*µ*_*V*_, *σ*_*V*_, *τ*_*V*_) as a function of (*v*_*E*_,*v*_*I*_,*w*) we evaluate the output firing rate of a neuron according to the following formula:

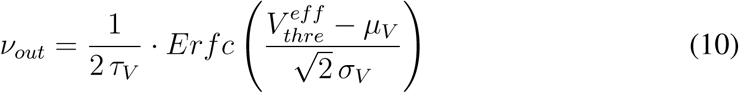

It has been shown, both theoretically and experimentally (Zerlaut et al., 2016), that the voltage effective threshold 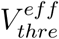 can be expressed as a function of (*µ*_*V*_, *σ*_*V*_, *τ* _*V*_). In particular, the phenomenological threshold was taken as a second order polynomial in the following way:

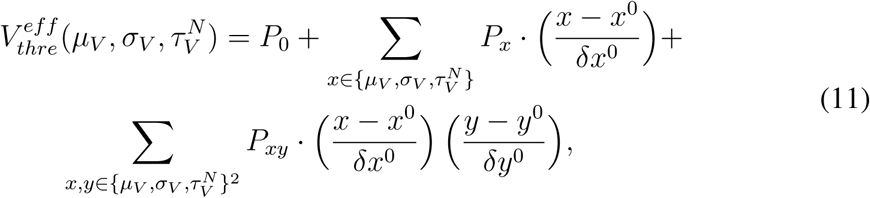

where we introduced the adimensional quantity 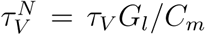. We evaluated {*P*} through a fit according to simulations on single neurons activity setting first 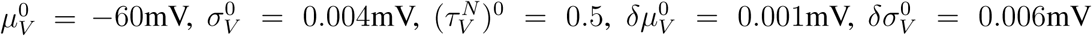 and 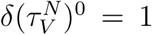. In general, the values of {*P*} do not depend on the parameters of single neuron dynamics (leakage, adaptation etc..) but we found a small dependence on the values of the neurons coupling parameters and on the parameters of the exponential term in single neuron dynamics that has been neglected in the derivation of membrane voltage moments. We perform the fit and calculate {*P*} in a case without adaptation for the sake of simplicity. We will use always this values of {*P*} for all the analyses performed in the paper (see table 1).

**Table 1:**
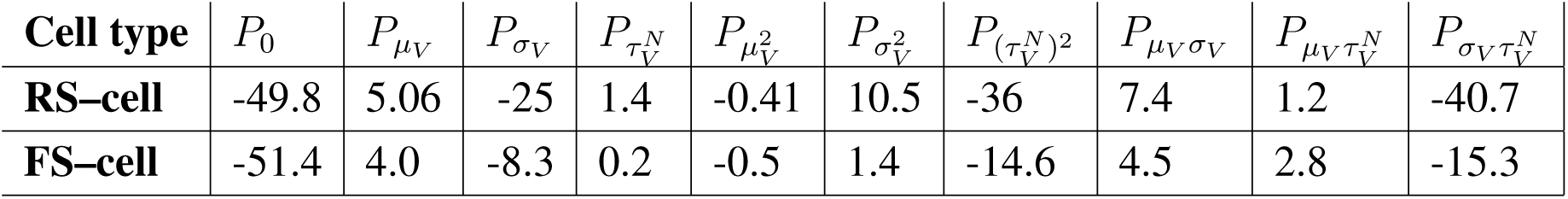
Fit parameters (expressed in mV)

The strong point of this analysis is that, once the fit is performed for a neuron with *b* = 0 and other fixed parameters, like the leakage currents, the same transfer function works also far from this fitting point because the effect of *b*, *E*_*L*_ and the other parameters is included in the theoretical evaluation of (*µ*_*V*_, *σ*_*V*_, *τ*_*V*_) whose values define the neuron output firing rate. In principle, the fit could be performed for any values of the parameters.

In Fig. 1 we show the result of the procedure here adopted for the evaluation of the transfer function. We consider RS-cell with a relatively high adaptation (*a* = 4, *b* = 20). We observe that this method is able to capture both neurons firing rate (upper panel) and their average voltage (lower panel). Moreover, we notice that not considering adaptation in the calculation of *µ*_*V*_ overestimates RS-cell voltage and would lead to a dramatic discordance for neuron firing rate (see black dashed line). In the methodology we use here we estimate the average voltage calculating adaptation in its stationary state, yielding the following result:

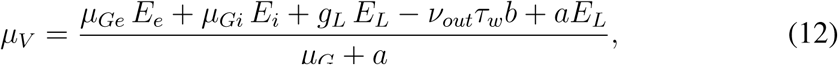

where *v*_*out*_ is the predicted firing rate of the neuron according to the transfer function.

**Figure 1:**
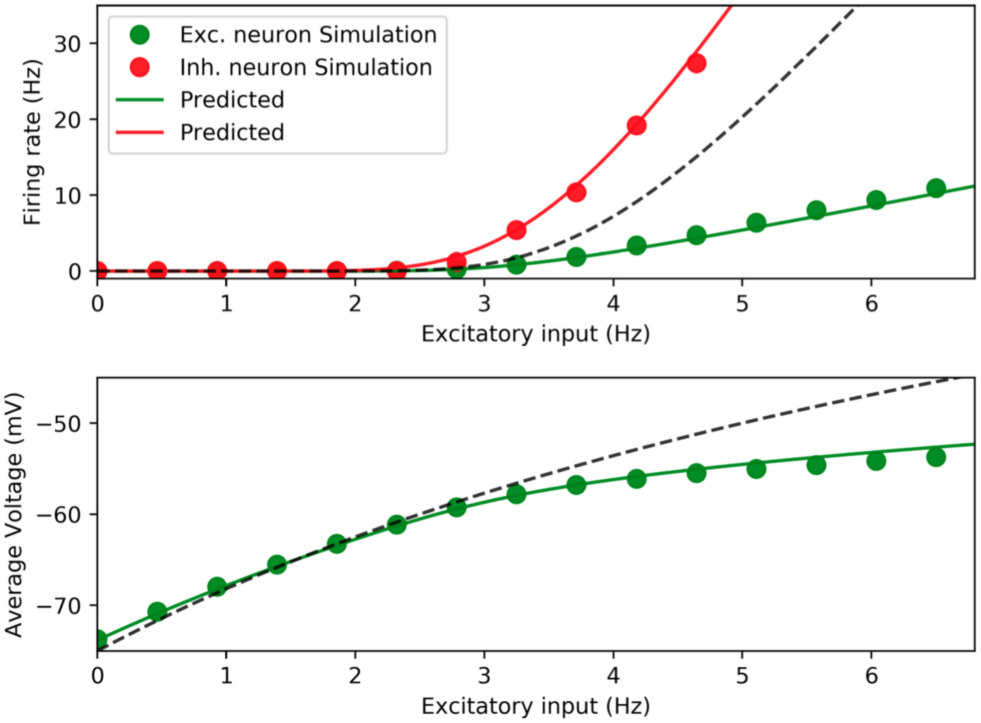
Transfer function evaluation. Top: RS (green) and FS (red) neuron stationary firing rate in function of the input excitatory firing rate *v*_*E*_ for *v*_*I*_ = 8*Hz*. Dots are direct simulation and continuous line prediction based on the semi-analytical transfer function. Black dashed line is the prediction obtained neglecting adaptation in the evaluation of neuron depolarization *µ*_*V*_ (see lower panel). Down: RS-cell average depolarization: dots direct simulation, line prediction based on Eq. (12) and dashed line prediction neglecting adaptation. Observe the necessity of a good evaluation of *µ*_*V*_ to predict correctly neuron firing rate (upper panel).

## 3 Results

Equipped with the formalism described in the previous section, we analyzed the dynamics produced by the mean-field model and compared them to the simulated neural network dynamics. We performed this analyses focusing on the response to various time-dependent input stimuli and the transition to oscillatory regimes by exploring different directions in the parameters space (adaptation strength, neurons excitability etc..).

### 3.1 Network spontaneous activity and mean-field prediction

We started by considering a region in the parameters space in which an excitatory external drive *v*_*drive*_*>* 0 is necessary in order to have spiking activity in the network. This corresponds, as we will see later, to choose low excitability level (low leakage reversal) for RS cells, i.e. 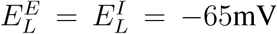. We chose *v*_*ext*_ = 2.5Hz, which guarantees an asynchronous irregular (AI) network dynamics with physiological values of conductances and neuron firing rates. As a first step we investigated the effect of adaptation on the spontaneous activity of the network, verifying the prediction capability of the meanfield model, focusing on the role of the adaptation strength *b*. We report in Fig. 2 the result of a network simulation, recording the average firing rate *v*_*e*_ and *v*_*i*_ of excitatory and inhibitory neurons (Fig. 2A) as well as the average voltage (averages are intended over time and over all neurons of the same type) *µ*_*V*_ (Fig. 2B). By increasing adaptation strength *b* we observe, as expected, a decrease in both excitatory and inhibitory firing rates (panel A, respectively green and red dots), accompanied by a decrease in *µ*_*V*_ (panel B, blue dots).

**Figure 2:**
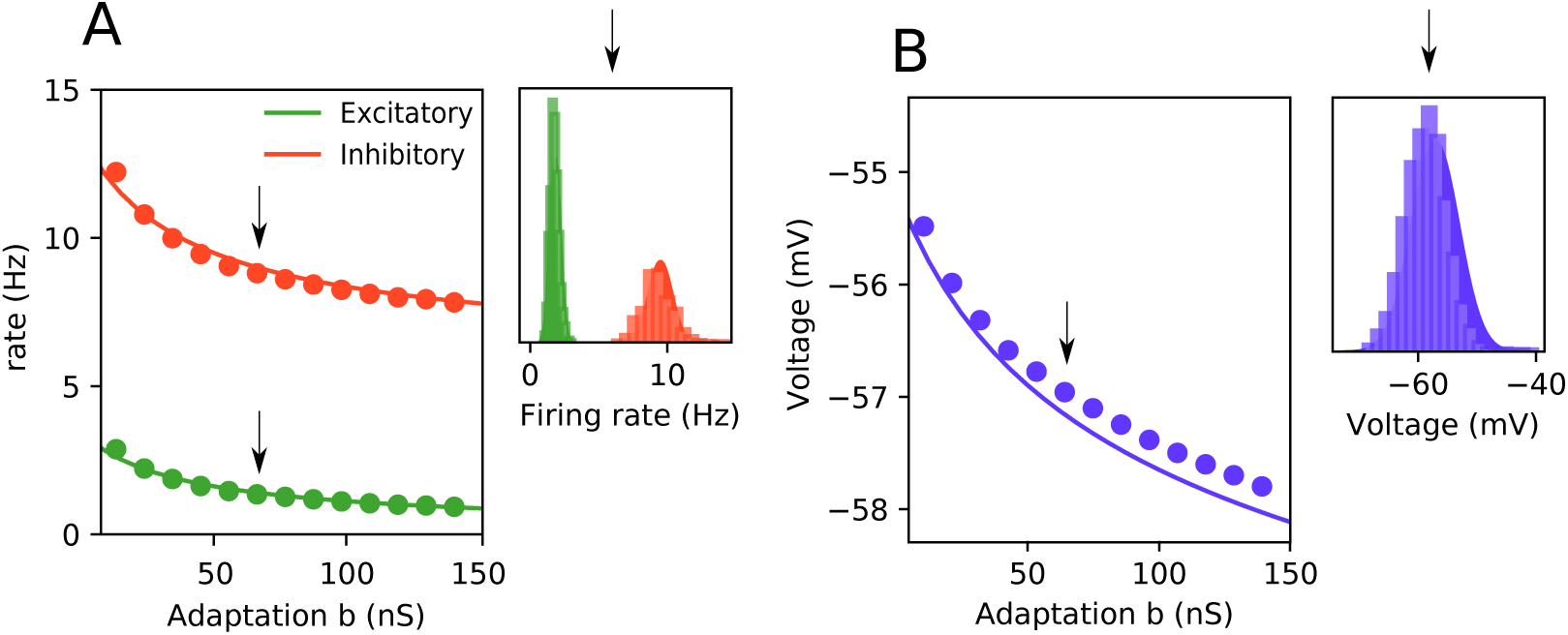
Spontaneous activity. **Panel A:** average firing rate from spiking network simulation. Solid lines are the means predicted by the mean-field model. In the right inset we report the firing rate distribution sampled from the spiking simulation (histogram) and the theoretically predicted Gaussian distribution (shading), for *b* = 60nS. Green and red are consistently referred to excitatory and inhibitory neurons. **Panel B** (Dots) average membrane potential from spiking simulation and (line) theoretical prediction. Inset, membrane potential distribution sampled from spiking simulation (histogram) and theoretically predicted distribution (shading), for the excitatory neurons for *b* = 60nS.

These quantities can be calculated from the mean-field model with adaptation described in Sec. 2. The average voltage *µ*_*V*_ is obtained from Eq. (12). In the same figure, we compare the simulation results with those of the mean-field model for the firing rates of the population (green and red solid lines in Fig. 2A for excitatory and inhibitory populations) and for the membrane potential (blue solid line in Fig. 2B). In the insets we report the comparison between the distribution of values in simulations and the theoretical results in the mean-field, for the specific parameter value *b* = 60nS. We observe that the mean-field is able to capture the spontaneous activity of the network and its fluctuations. It is worth notice that the spontaneous activity of the network could have been captured also with a ‘naive’ version of this mean-field model considering only stationary values of adaptation and not its time dynamics (for the sake of simplicity we refer to this version of the mean-field model as “stationary” or “non–adaptive”).

### 3.2 Response to external stimuli

Even if the spontaneous activity of the populations firing rate in the network could be sufficiently well predicted also by using a “stationary” mean-field, in this section we will show how adaptation dynamics needs to be taken into account when the stationarity condition is not satisfied^1^. In particular we studied the response of the network to time-dependent external stimuli. We report in Fig. 3 (A1-A3) three different examples of external stimuli:

- In Fig 3-A1 we add an extra stimulus *v*_*ext*_ with exponential rise and decay, to the external drive (see dashed line in lower inset). If the rise and decay time scales are smaller than the time scale of the adaptation dynamics (as it is usually the case, since adaptation is a slow variable – *τ* _*w*_ = 500ms in our case) only a meanfield model taking adaptation *W* into account can predict the network response in a correct way. By looking at the comparison between a mean-field model with or without adaptation (green vs. dashed blue line) we observe that the extended version of the mean-field correctly captures the first peak of response, as well as an hyper-polarization at the stimulus offset due to the accumulation of adaptation.
- In Fig. 3-A2 we consider a input yielding an inhibition by temporarily turning off the initially constant external drive *v*_*drive*_. We observe a rebound response captured very well in its time course by the mean-field model. The effect of adaptation is therefore very strong also by looking at the network response after inhibition.
- In Fig 3-A3 we show the response to an oscillatory input of a fixed frequency *f*_*inp*_. It is evident how the effect of adaptation allows the mean-field to catch the time-dependent amplitude of the response, while it is always constant and underestimated in the naive case of “stationary” mean-field. Adaptation dynamics is then responsible for network increased response to an oscillating external input *v*_*ext*_.

**Figure 3:**
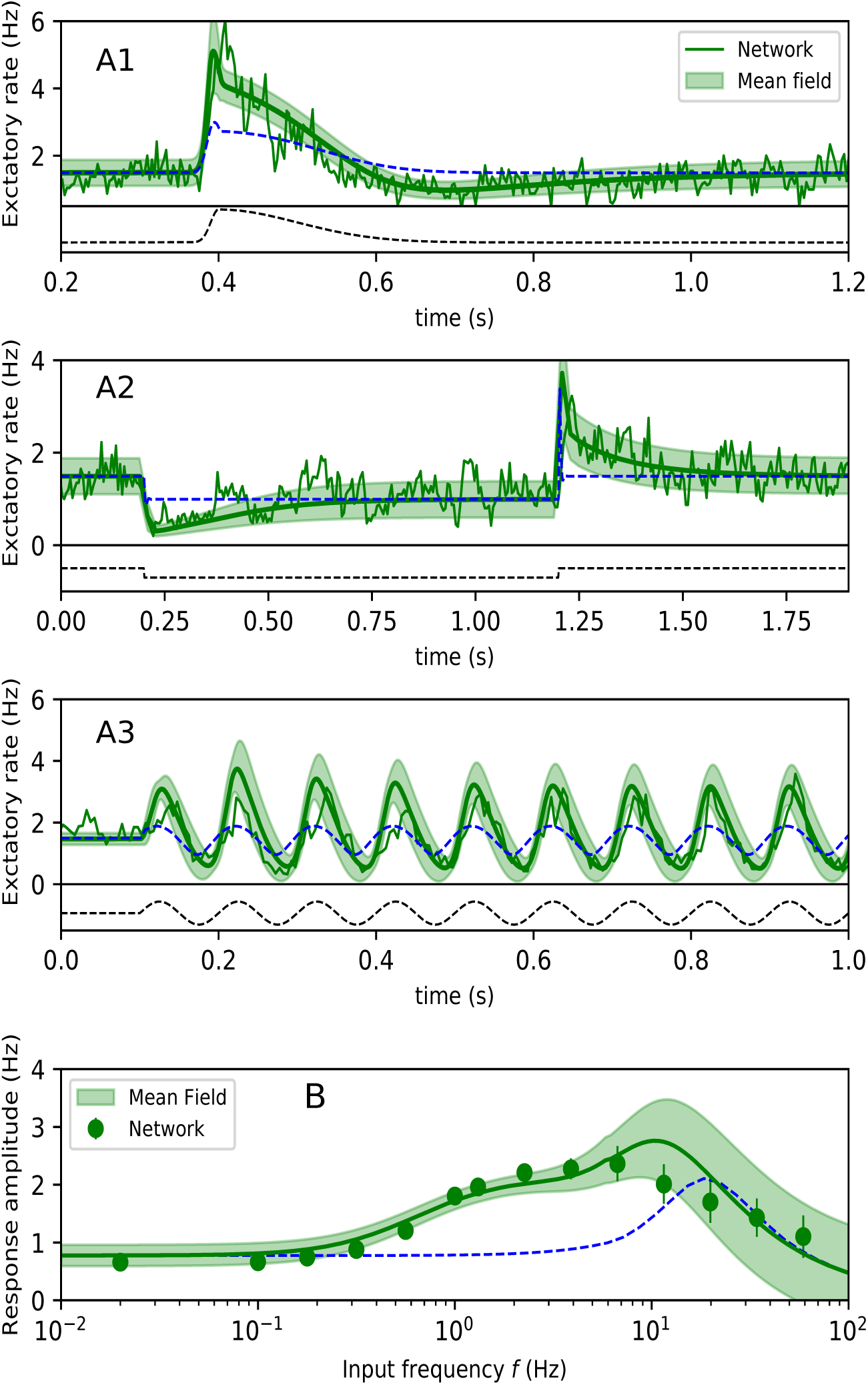
Response to external stimuli. **A1-3** Population activity of the excitatory sub-populations in presence of different time-varying external stimuli (green). Superimposed is the mean and standard deviation over time predicted by the Markovian formalism. (bottom) Time-course of the external stimulus. (blue) The theoretical prediction when the adaptation variable *W* is fixed to its stationary value 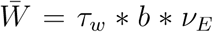. **B** (Dots) Amplitude of network oscillations, in response to an oscillating input as a function of the input frequency. Superimposed is the mean and standard deviation predicted by the model. (blue) The theoretical prediction when the adaptation variable *W* is fixed to its stationary value 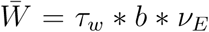. The parameter *a* is set to zero in these simulations.

In Fig. 3D we study more extensively the response of the network to oscillating external inputs, as a function of *f*_*inp*_. In particular, we show that the amplitude of the oscillations of the firing rate (the difference between the maximum and the minimum firing rate once the system has passed the transient phase) has a very non-trivial behavior when the frequency varies. First, there is an increase at *f*_*inp*_ ∼ 2Hz. Then, a peak appears around 10-50 Hz (Zerlaut et al., 2018). Finally, a maximum peak is followed by a drop at a second time scale around *f*_*inp*_ ∼ 20Hz. The mean-field is able to predict these combined effects and gives us an indication on their origin. The rise observed at *f*_*inp*_ ∼ 0.2Hz can be related to the time scale of adaptation *τ*_*w*_ ∼ 1*/f*_*inp*_. In fact, without adaptation, the mean-field (dashed blue line of Fig. 3D) does not show the same increase at low frequencies and instead it is completely transparent to the frequency of the external input, until the appearance of a resonance peak. It can be understood given the relatively high strength (compared to the external input) of the excitatory-inhibitory loop, bringing the system close to a bifurcation toward oscillations ((Brunel and Wang, 2003)) and when forced at the correct frequency, the response is amplified ((Ledoux and Brunel, 2011)). This is the case for both mean field models, with or without adaptation. Finally, the decay from the baseline response amplitude appears at frequencies of order 1*/T* and can be easily understood by observing that when the stimulus varies faster than the correlation time scale of the mean-field, it appears as an effective constant external drive and the oscillations disappear. Consistently, the same decay at high frequencies is conserved in the “stationary” mean-field.

### 3.3 State-dependent responsiveness

We study in this section the response of the network as a function of its dynamical state preceding the input arrival.

The input here considered corresponds to one cycle of a sinusoidal wave of spike train at a frequency *f* = 5*Hz* (see insets in Fig. 4). We study two different parameter setups that differ for the baseline drive the system receives. In case (A) *v*_*drive*_ = 7*Hz* and the system sets in an asynchronous state with relatively high firing rate and very high conductance level (*G*_*E*_*/G*_*l*_ ∼ 3) while in case (B) *v*_*drive*_ = 1.5*Hz* and neurons firing rates are lower and conductance state has realistic values (*G*_*E*_*/G*_*l*_ ∼ 0.8). We observe that, for the same stimuli, network (B) has a much greater response to the input with respect to network (A). Moreover, the mean field model is able to capture this difference and gives a very good prediction of response time course. This effect is even stronger when comparing the relative response of the two networks with respect to their baseline (see in Fig. 4C the comparison between the two continuous lines). The state-dependent responsiveness of the system is a combination of two effects: the dynamics of adaptation and the conductance state. In order to elucidate this mechanism we report in Fig. 4C (dashed lines) the responses in the “stationary” model, as done in Fig 3. We observe that such model does not capture the right peak in response to the stimuli. In fact, the peak is strongly affected by the level of adaptation pre-stimulus, which is quite low as the excitatory neurons firing rate is low. Accordingly, when a sufficiently fast stimuli (with respect to the time scale of adaptation, as it is the case for 5Hz) is presented, the system will strongly activate. The dynamics of adaptation is thus responsible for a good part of the state dependent response, due to the lower or higher pre-stimuli adaptation/excitatory neurons firing rate. On the other hand, also a model that does not take into account the time evolution dynamics of adaptation (dashed line) shows an increased responsiveness in the lower conductance state. This shows the importance of using a mean-field model taking into account both conductances and adaptation dynamics.

**Figure 4:**
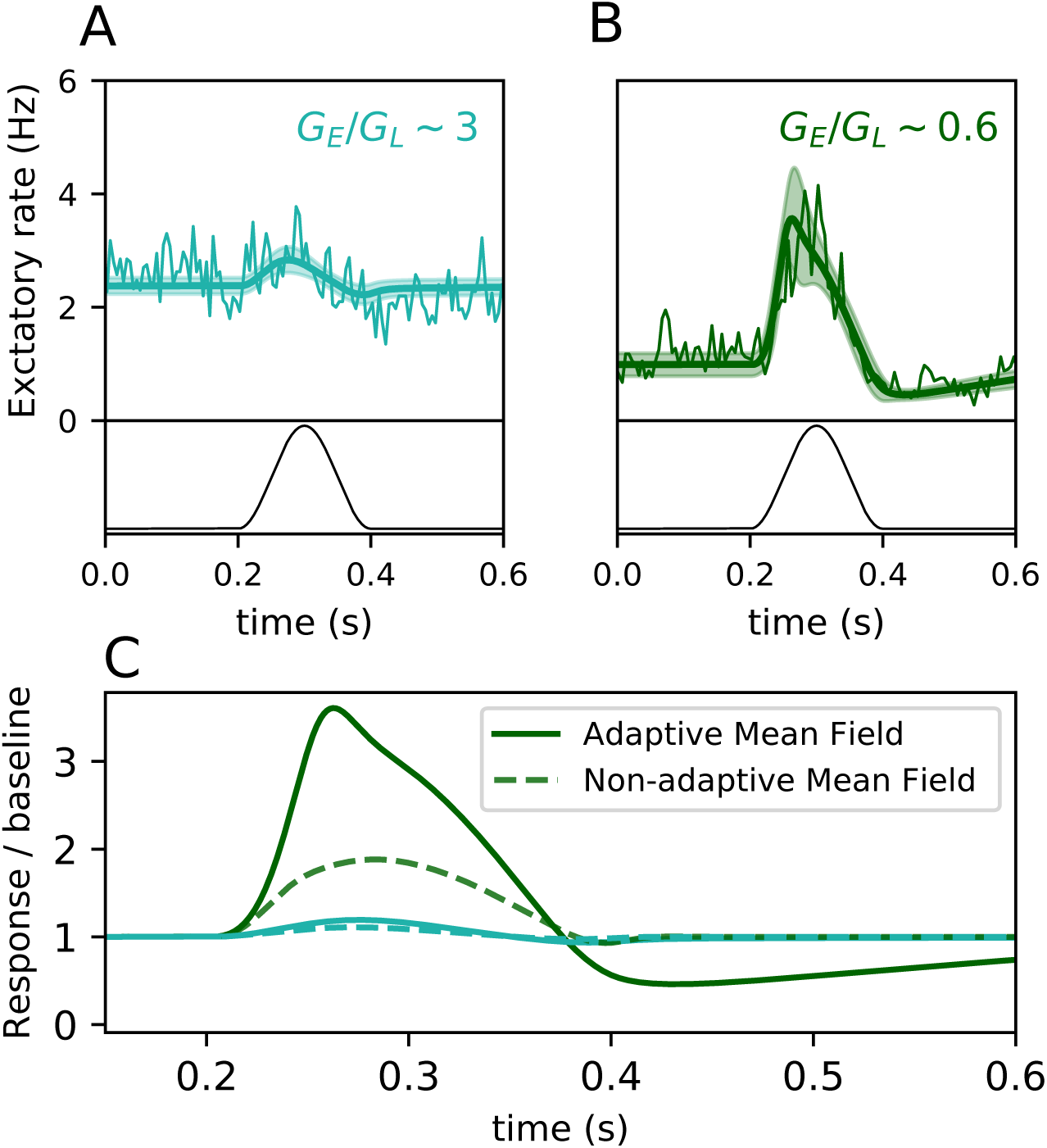
State-dependent response. Population activity of the excitatory subpopulations in response to a cycle of a sinusoidal input of amplitude 2Hz (see lower insets). Superimposed is the mean and standard deviation over time predicted by the Markovian formalism. Green dark color (B panel) has a different baseline firing rate with respect to light cyan (A panel) due a different *v*_*drive*_ (*v*_*drive*_ = 1.5Hz for green and *v*_*drive*_ = 7Hz for cyan). The dashed line correspond to the mean field model where adaptation is not evolving in time but takes values corresponding to its stationary value (as in Fig 3).

### 3.4 Transition to self-sustained bistable network activity

In the setup used so far, the network needs a constant external drive *v*_*drive*_ in order to be set in an AI state. In the absence of external drive, *v*_*drive*_ = 0, the only stable state of the system is silent, with *v*_*e*_ = *v*_*I*_ = 0. This is true for the case we investigate here of adaptation set to zero. We will reintroduce adaptation in next section to study its effects.

We show in Fig.5 that it is possible to observe a transition to a bistable network, by modifying the excitability of RS and FS cells. This is done by changing the resting potential (actually changing the leakage reversal potentials 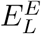 and 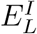). In order to verify the existence of a bistable network dynamics in the spiking network we perform a simulation with an initial kick (*v*_*drive*_ = 1 Hz) applied to the network for a small period of time (around 100 ms). In the case of a bistable system the network activity rises and then remains to a non-zero firing rates, even after the end of the stimulus. On the contrary, the system will come back at some point to a silent state. We report in the inset of Fig.5A the two cases (i) 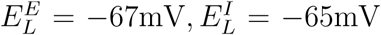 (red triangle) and (ii) 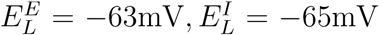 (green square). By plotting the firing rates average after the stimulus offset (we measure for 5s starting 1s after the stimulus offset) as a function of 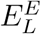 and 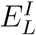, we observe the existence of a clear transition line separating a bistable regime to a state with only one silent stable state Fig.5A. The bi-stability regime is achieved as soon as the excitability of RS cells overcomes the excitability of FS cells by a certain amount. In particular, the transition line (the couples of critical points 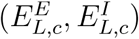) lies almost at the bisector of the square, meaning that the critical point is achieved at the critical ratio 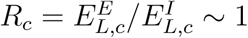.

The existence of two stable fixed points in the network dynamics can be then investigated in the mean-field model. We report in Fig5B a graphical solution for the fixed point of *v*_*e*_. It is calculated as follows. We scan the values of *v*_*e*_ (x-axes) and we calculate the corresponding 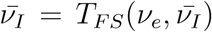 (1st order mean-field 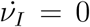). We then calculate the corresponding 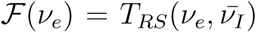. We obtain a fixed point of the system when ℱ(*v*_*e*_) = *v*_*e*_. We thus plot the function ℱ and search for intersection with the bisector. Let us consider the case (i) (see heat plot in Fig5A) which corresponds to parameters for which in the network we observe only one silent stable state. We observe that actually the mean-field predicts correctly that there is only one intersection at *v*_*e*_ = 0. Let us notice, for the sake of completeness, that it exists always a couple of other fixed point not shown in the plot at very high frequencies (one stable and one unstable), corresponding to the unrealistic case *v*_*e*_ = *v*_*I*_ = 1*/T*_*refr*_. Increasing the excitability of RS cells (case (ii)) we observe the appearance of two new fixed points. We verified that the higher in firing rates is indeed stable. We then superimpose the firing rate of the network after a kick for these values of the leakage (green dots), and we observe that actually the mean field fixed point matches with network simulations. Using this method we are able to calculate the transition curve from a self-sustained to a non-self-sustained regime in function of 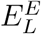 and 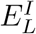, that we superimpose in the spiking network simulations reported in the heat-plot of Fig. 5A, finding a very good agreement.

**Figure 5:**
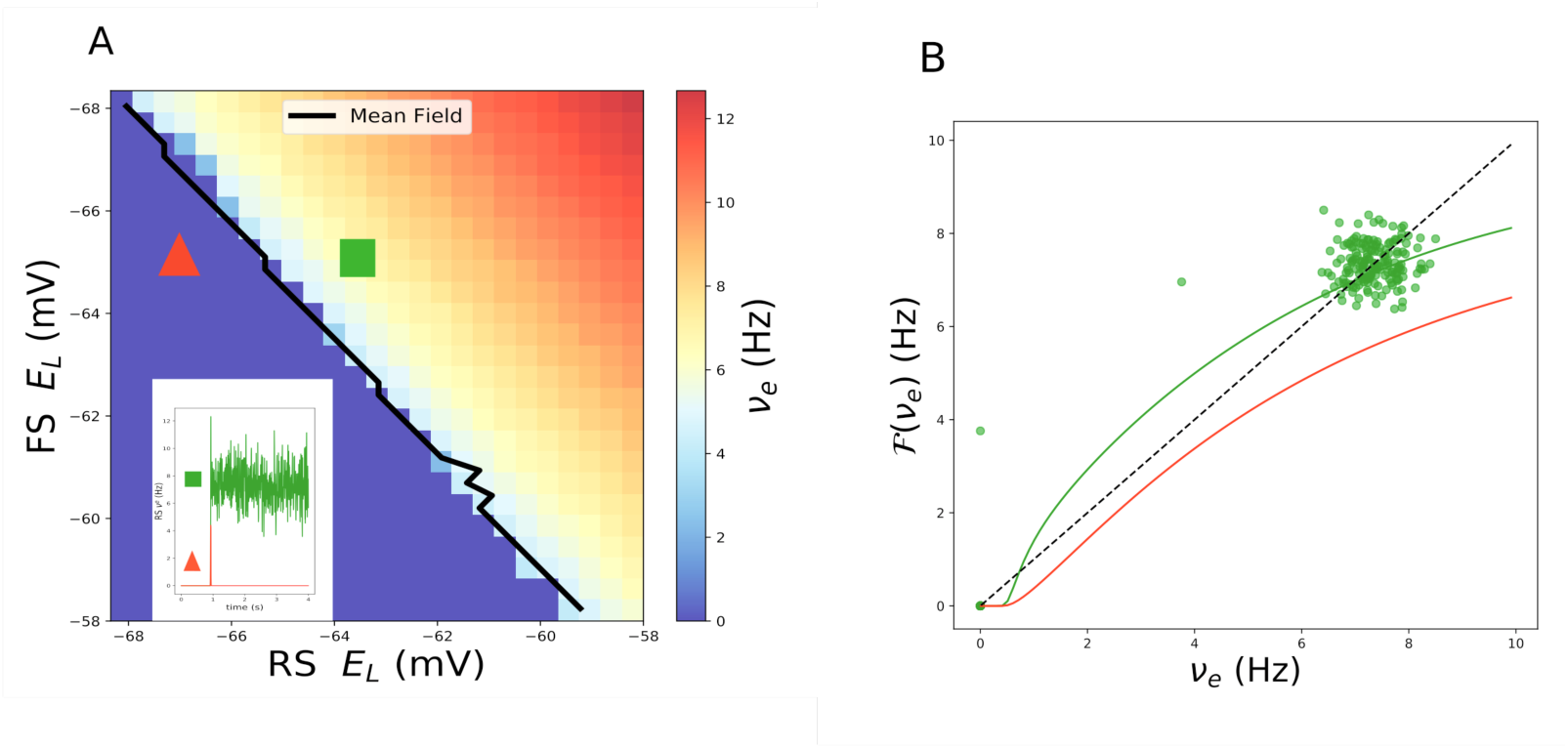
Transition to self-sustained activity. A: Firing rate of excitatory neurons (averaged over 5s) after the kick of duration of 100ms. The black line is the prediction from mean field for the transition point. In the inset the excitatory response of the network to the kick in two cases 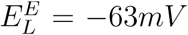 and 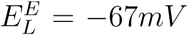. B: Function ℱ derived by mean-field equations in the two cases (green and red curves). The dashed black line is the bisector. Green light dots have been obtained from the network simulation shown in the inset of panel A. Adaptation is set to zero in this simulation.

Let us notice that, again, even if the fit for the transfer function (see methods) has been done for a specific value of 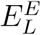 and 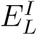, it still gives correct results also moving around in the parameter space.

Moreover, we point out that this self-sustained regime is characterized by physiological conductances, observing a quite large range of parameter values where the conductances stay at a physiological value, at variance with other models present in the literature ((Vogels and Abbott, 2005)).

### 3.5 UP-DOWN states dynamics triggered by noise and adaptation

We report here the effects of adaptation on the network dynamics in the case of a bistable system (case (ii) of the previous section).

In the absence of any external drive the system has still the same dynamics as in the case without adaptation, i.e. silent or self-sustained state. Nevertheless, this state is less stable the more the adaptation strength increases. We consider *b* = 60 nS, yielding a realistic level of adaptation to RS cells. For this parameter value the active state is unstable and only a silent state is permitted.

As soon as a small external drive is added to the system (here we use *v*_*drive*_ = 0.315 Hz), it introduces a noisy level of activity in the silent (down) state. As in the down state adaptation is almost zero, the second “active” state is stable. Accordingly, noise permits to the system to “jump” to the active (UP) state. Nevertheless, as adaptation grows (because of neurons firing), the UP state loses stability and the network goes back to the DOWN state (see Fig. 6). The duration of the UP state is related to the speed at which adaptation grows as function of the neurons firing rate increase. Nevertheless, we observe that UP state durations are heterogeneous and characterized by “bumps” of activity, revealing a non-trivial dynamical structure that is induced by finite size noise fluctuations. The alternation of UP states is irregular, as their duration and structure (see Fig. 6A-B).

**Figure 6:**
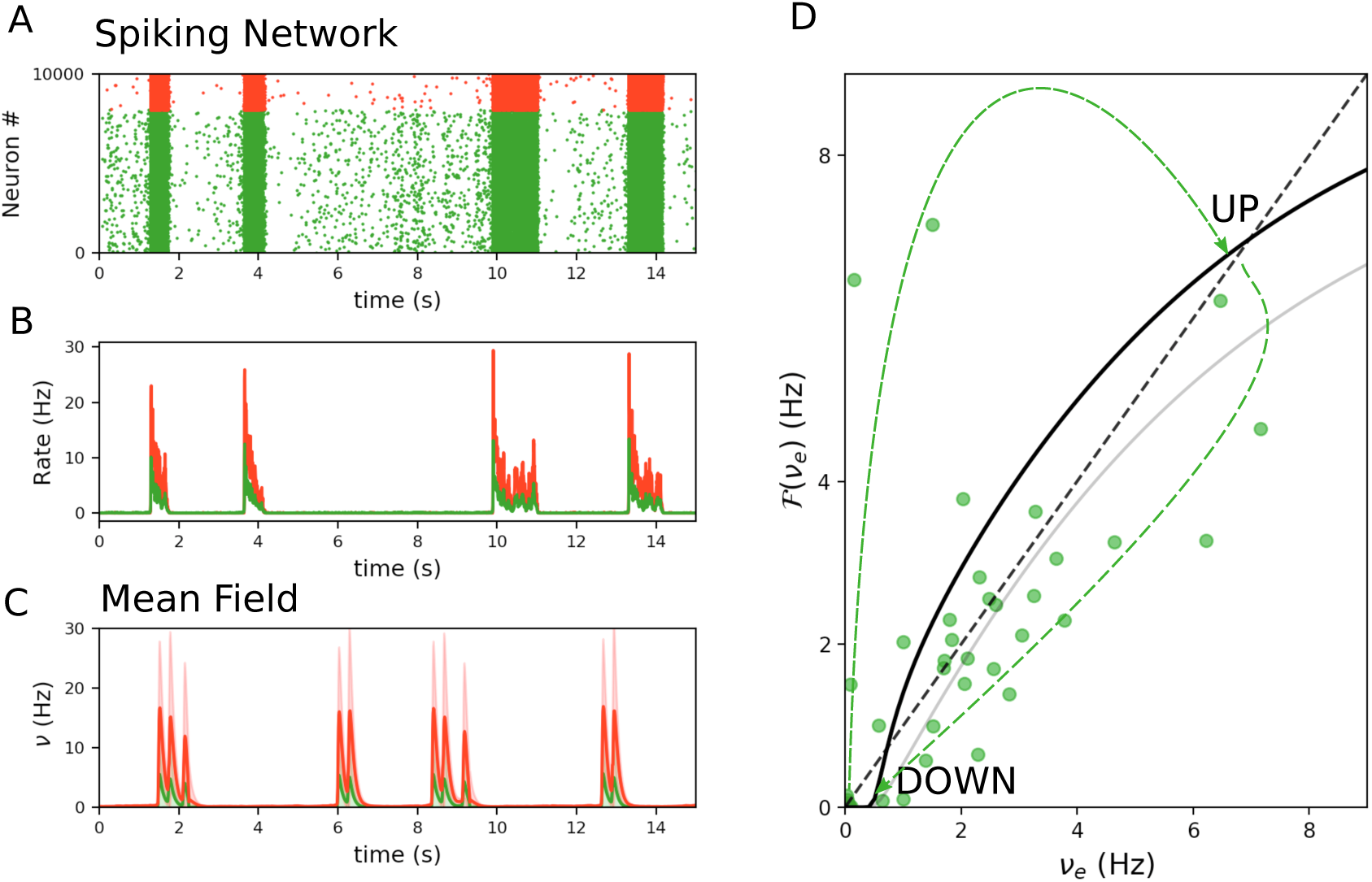
UP DOWN state dynamics in bistable network. Dynamics of a bistable network (RS *E*_*L*_ = 63*mV*) with an external Poissonian input *v*_*ext*_ = 0.315 and adaptation *b* = 60pA. A-B Raster plot and average firing rate in the spiking network (green stands for excitation and red for inhibition). C Corresponding mean field model dynamics with external additive noise (see text). D Phase plane derived from mean field with superimposed the firing rate (green dots) of the network dynamics (panel B). Black line is relative to the transfer function for *E*_*L*_ = 63*mV* (zero adaptation) and grey line is relative to a lower leakage reversal due to adaptation building up during the UP state. Green dashed arrows are used to guide the eyes through the spiking network trajectory during DOWN-UP cycle.

The deterministic mean-field model with adaptation used until now cannot reproduce this kind of dynamic as it does not take into account the amount of noise induced by non-zero external drive (*v*_*drive*_ = 0.315 Hz). In order to account for noise induced by an external drive *v*_*drive*_ of Poissonian spike trains targeting both excitatory and inhibitory networks we consider an additive noise modeled as an Ornstein–Ulhenbeck (OU) process to Eq.s 4. Accordingly, the external drive *v*_*drive*_ becomes here a time dependent variable *v*_*drive*_(*t*) = *v*_*drive*_ + *σ ξ*(*t*), where *ξ*(*t*) is an (OU) process with a fast time scale of 5ms, σ = 10.5 is the noise amplitude and *v*_*drive*_ is the average of the process *v*_*drive*_(*t*).

A simulation of the mean-field model in this set-up is reported in Fig.6C. We observe the alternation of silent periods with transients of high activity. The firing rates during the UP states are quantitatively matching those of the spiking network simulations, as their duration Finally, a certain level of heterogeneity in between UP states is reproduced by the mean field, where rebounds of activity are present, exactly as in spiking network simulations. In Fig.6D we superimpose the network dynamics during an UP-DOWN cycle to the activity map derived for the mean field (like done in Fig5B). We observe that the trajectory drawn by the network follows the stability principles dictated by the mean field. The system is dynamically bistable when adaptation in zero (DOWN state) and can jump to the UP state. Then, when adaptation builds up, the system is not bistable anymore and comes back to the DOWN state.

## Conclusions

In the present paper, we derived a “biologically realistic” mean-field model of neuronal populations, that includes nonlinear effects important for neural dynamics. Our approach was similar to a previous Master Equation formalism (El Boustani and Destexhe, 2009), which we have augmented by explicitly including the dynamics of adaptation. We discuss this model, how it relates to previous approaches, and what perspectives it opens for future work.

The main originality of the mean-field model proposed here is that it takes into account the presence of strong nonlinear effects such as conductance-based synaptic interactions, and spike-frequency adaptation. To do this, we went back to first principles and re-derived the previous Master Equation formalism to take into account slow variables like adaptation, taking into account the “memory” of the network dynamics, which was not considered in the original Markovian formulation (El Boustani and Destexhe, 2009). The TF of neurons is obtained using a semi-analytic approach, as done previously (Zerlaut et al., 2016). This allowed us to obtain a mean-field model for networks of spiking neurons, where excitatory and inhibitory neurons have different intrinsic properties (RS and FS cells), with conductance-based synaptic interactions and where adaptation is taken into account.

The mean-field model was tested by comparing its predictions to the full spiking network model, and it was found that several properties are correctly captured. First, the model correctly predicts the level of spontaneous activity in AI states. Networks of RS and FS cells are characterized by AI states where RS and FS cells display different levels of spontaneous firing, with higher frequencies for FS cells. These features are observed experimentally in cortex, where inhibitory neurons have a higher level of firing (Roxin et al., 2011; Dehghani et al., 2016). The present mean-field model predicts the level of firing activity when adaptation has settled to a steady-state, which was not possible in previous mean-field models of AdEx networks (Zerlaut et al., 2018). In fact, if adaptation is not considered, an ad-hoc fitting is necessary to adjust the transfer function, making the all procedure satisfactory but limited to the fitting point (Zerlaut et al., 2018). The approach proposed here permits to avoid this problem and obtain a mean field model which stays valid even far from such fitting point, a necessary ingredient when we investigate the phase space of the model (see Fig. 5A) or the emergence of slow oscillations. In previous mean-field models, the level of spontaneous activity of IF networks with conductance-based synapses could be predicted (El Boustani and Destexhe, 2009), but this prediction was not quantitative because the transfer function was approximated (and of course no adaptation was present).

A second property is that the present mean-field model captures the full time course of the response of the network to external input. This includes the transient initial response, the peak of the response and the “tail” at longer times, where adaptation plays a role. Previous mean-field models could predict the response dynamics of networks, but only for current-based synaptic interactions (Schwalger et al., 2017; Montbrió et al., 2015). The present mean-field model captures the response of conductance-based spiking networks remarkably well, including complex stimuli like oscillations at different frequencies. The whole spectrum of oscillatory responses could be well predicted (Fig. 3–4), while previous mean-field models of AdEx networks typically failed to capture the frequency response (Zerlaut et al., 2018). This constitutes a major improvement, but most importantly, it suggests that the present mean-field model should be able to adequately capture the dynamics of interconnected networks, which opens the perspective of more realistic modeling of large-scale systems. This constitutes an exciting perspective for future work.

A third important feature reproduced by this model is that the same network can produce different responses to the same input, according to its level of spontaneous activity. Such state-dependent responses are found experimentally at various levels, from single cell level (Haider et al., 2007; Hasenstaub et al., 2007; Sachdev et al., 2004; Timofeev et al., 1996; Reig and Sanchez-Vives, 2007; Reig et al., 2015; Shu et al., 2003) to networks and large-scale systems (Silvanto et al., 2008, 2007). In the model, we found that such a state dependency is due to the fact that different levels of activity will set neurons in different conductance states, and thus individual neurons will have different responsiveness. The steady level of adaptation is also dependent on the level of spontaneous activity, and also contributes to the state-dependent response. To our knowledge, no previous mean-field model is able to display such state dependency, and this constitutes a significant advance in the biological realism of mean-field models.

A last important property reproduced by the mean-field is that networks of neurons with adaptation can produce UP/DOWN state dynamics (Steriade et al., 2001; Timofeev et al., 2000; Compte et al., 2003). Although simplified models were also proposed for UP-DOWN state oscillations (Capone and Mattia, 2017; Jercog et al., 2017), there is at present no mean-field model of such adaptation dynamics derived from conductance based spiking networks. If used in a larger-scale system, the present mean-field model should be able to reproduce the dynamics of slow-wave activity at larger scales. This was never done using realistic mean-field models and this also constitutes a possible extension of the present work.

In addition to those listed above, several additional possible extensions of the present model could be considered. First is to extend to formalism beyond spike-frequency adaptation. A similar approach could potentially be used for other purposes, e.g. other external variables like Spike-Time-Dependent-Plasticity (STDP), capable to yield different dynamical regimes with respect to those investigated here (Tsodyks et al., 1998) or synaptic dynamics in general. In fact, a possible extension of this model is to consider fast oscillations, typically due to synaptic or delay dynamics of excitation-inhibition (Buzsáki and Wang, 2012; Bos et al., 2016). In our mean field model the dynamics of synapses is not considered dynamically but a possible extension, using the same protocol of the one here introduced for adaptation, might be performed. Moreover, the approach we used to calculate the transfer function is very general and it may be applied to other neuronal models or to real data (Zerlaut et al., 2016), provided the neuron dynamics has a stationary firing rate. In fact, if neurons display mechanism like bursting (Izhikevich, 2003) the calculation of the stationary transfer function can be not even well defined. For instance, it has been shown that neurons may display stochastic resonance or in general, a non-trivial response in the frequency domain (Lindner and Schimansky-Geier, 2001). For these classes of neurons a different approach should be implemented, calculating the transfer function in the frequency domain. Nevertheless, for the cortical regimes we described here, with a high realistic features, our approach was very satisfactory.

Another possible extension is to include the heterogeneity of the TF of neurons found experimentally in mouse cortex (Pospischil et al., 2008; Zerlaut et al., 2016), where the parameters of adaptation dynamics strongly vary across neurons. In the present mean field model formulation, neuronal heterogeneity is not taken into account and might represent a future development of this work, similar to previous work (di Volo et al., 2014), in order to obtain a heterogeneous mean field model based on experimental measures (Zerlaut et al., 2016).

Finally, another extension is to further explore the state-dependent responses, in the context of the detection of external stimuli or sensory awareness. This could link the present approach to modeling different levels of sensory awareness in large-scale multi-areal networks. We believe that such models are relevant to biological data only if they include biologically relevant features like state-dependent responsiveness. The present mean-field model is to our knowledge the first one to account for such state dependency, and thus should be considered as a step towards building biologically-realistic large-scale models at mesoscopic scale.

## Acknowledgements

We acknowledge funding from the European Union (Human Brain Project H2020– 720270 and H2020–785907). We thank Yann Zerlaut for useful discussions.

## Appendix Master equation formulation in presence of slow variable.

We extend here the framework discussed in (El Boustani and Destexhe, 2009) using a consistent notations. Let us consider a network with *K* homogeneous populations of neurons. Each population is *γ* defined by its network activity *m*_*γ*_ that is the number of neurons which fired in that population in a time bin *T*. In the wit of making a probabilistic Markovian formulation, *T* should be chosen of the time-scale of correlation decay in order to have the system that only depends on the previous step. Also *T* should be small enough to avoid to have the same neuron firing twice in the same bin.

We also define the variable *W*_*γ*_ = (1*/N*_*γ*_) ∑_*i*_*w* _*γ,i*_, where *w*_*γ,i*_ is the adaptation of the *i – i th* neuron in population *γ*, as defined in the previous paragraph. The dynamics of the variables *W* is assumed to be slow with respect to the autocorrelation time of the system *T*.

We make the assumption that the state of the system is defined by the set of variables {*m* _*γ*_, *W* _*γ*_ }. The network behavior can be investigated by studying the transition probability 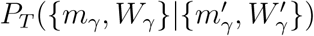, i.e. the probability that the system is in {*m*_*γ*_, *W*_*γ*_ } at time *t*_0_ + *T* conditioned to the fact that it was at 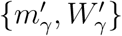 at a generic time *t*_0_. Provided the choice for *T* we discussed above, we can reasonably assume that population-conditional probabilities are independent beyond the time scale of T, thus allowing to write:

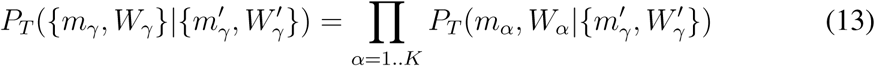

We can thus define the transition operator *W* as:

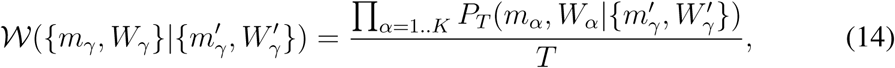

For this approximation to be valid, the time constant *τ*_*w*_*≫ T*, namely we assumed that the adaptation dynamics is slower than the firing rate dynamics.

This also involves that *W* variables are independent on fluctuations in firing rates and can be described by a deterministic equation. Thus the probabilities can be factorized as follows

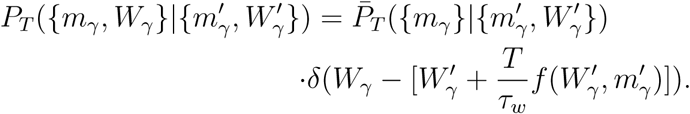

where we used Eulero integration for *W* dynamics that for linear *f*() can be explicitly written in the following closed form

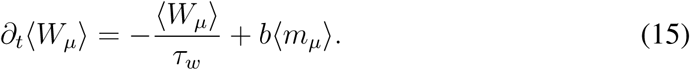

Notice that here for the sake of simplicity we consider *a* = 0 (see Eq. 1), neglecting voltage-dependent adaptation. The extension to *a* ≠ 0 is trivial once the average population voltage is calculated and is described in the model section.

Using the same approach as in (El Boustani and Destexhe, 2009) we obtain the following equations for the average activity and for the correlations

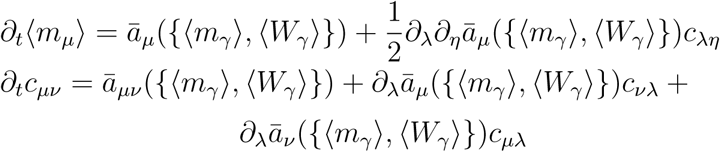

where

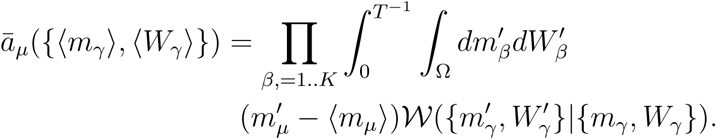

and

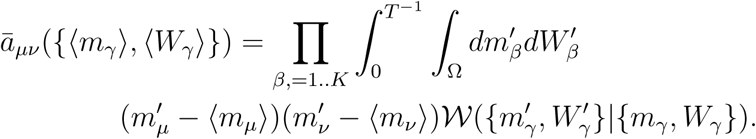

Using the assumption made in eq.(15) W can be explicitly written as

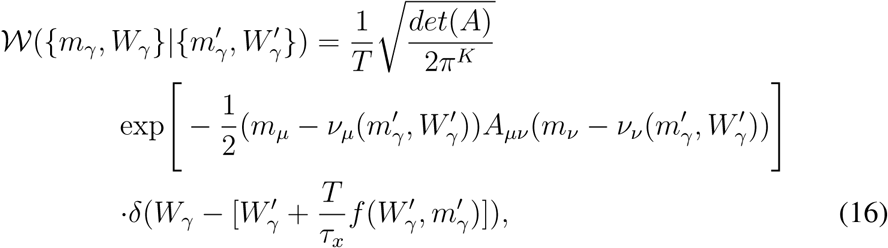

where *v*_*µ*_ depends on the single neuron model and where 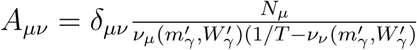. We finally get the equations for the moments:

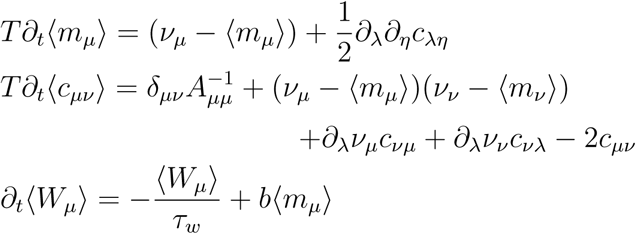

where, once again, only the first order for the equation of *W* are considered, since we suppose its dynamics not strongly affected by fluctuations. Here we stress that the activity variables dynamics are a function of the adaptation level.

## Footnotes

1 Notice that in the case of time-dependent external stimuli, the value of adaptation in the “stationary” mean-field is not constant. Nonetheless, it is “stationary” in the sense that at time *t* we assigned to it the value the adaptation would have in a stationary state with firing rate for the system as at time *t*.

